# The combination of CD49b and CD229 reveals a subset of multipotent progenitors with short-term activity within the hematopoietic stem cell compartment

**DOI:** 10.1101/2023.03.20.533430

**Authors:** Ece Somuncular, Tsu-Yi Su, Özge Dumral, Anne-Sofie Johansson, Sidinh Luc

## Abstract

Hematopoiesis is maintained by hematopoietic stem cells (HSCs) that replenish all blood lineages throughout life. It is well-established that the HSC pool is functionally heterogeneous consisting of cells differing in longevity, self-renewal ability, cell proliferation, and lineage differentiation. Although HSCs can be identified through the Lineage^−^Sca-1^+^c-Kit^+^CD48^−^CD34^−^ CD150^+^ immunophenotype, the cell surface marker combination does not permit absolute purification of functional HSCs with long-term reconstituting ability. Therefore, prospective isolation of long-term HSCs is crucial for mechanistic understanding of the biological functions of HSCs and for resolving functional heterogeneity within the HSC population. Here, we show that the combination of CD229 and CD49b cell surface markers within the phenotypic HSC compartment identifies a subset of multipotent progenitor cells with high proliferative activity and short-term reconstituting ability. Thus, the addition of CD229 and CD49b to conventional HSC markers permits prospective isolation of functional HSCs by distinguishing MPPs in the HSC compartment.

**SIGNIFICANCE STATEMENT:** Blood cells are generated by rare stem cells, which based on their defining features of self-renewal and multipotent abilities, are used in transplantation as curative treatment for blood diseases. Despite their clinical importance, mechanisms of stem cell properties are elusive given the inability to isolate pure blood stem cells for prospective analyses. In this study, we demonstrate that stem cells can be isolated by excluding multipotent progenitors, marked by CD229 and CD49b, from the stem cell compartment. Our study has important implications for the analysis of mechanisms regulating stem cell function and is relevant for clinical applications of stem cells.

## INTRODUCTION

Blood cells in the hematopoietic system are replenished by infrequent hematopoietic stem cells (HSCs). The hallmarks of HSCs are multipotency and self-renewal ability.^1,2^ Although HSCs can be identified by immunophenotype, they are only distinguishable from multipotent progenitors (MPPs) with short-term (ST) self-renewal potential through functional studies. Hematopoietic stem cells are defined by their ability to continuously repopulate short-lived mature blood cells in myeloablated mice for at least 4-6 months, with the capacity to reconstitute secondary transplanted animals.^2^

The HSC population is functionally heterogeneous, however, the mechanism for distinct HSC behaviors remains obscure.^3–11^ Hematopoietic stem cells reside within the phenotypically defined Lineage^−^Sca-1^+^c-Kit^+^ (LSK) compartment and can be enriched through CD34^−^,^12^ Flt-3^−^

,^13^ CD48^−^, and CD150^+^,^14^ cell surface marker selections. Inclusion of CD49b^15^ or CD229^16^ have been suggested to resolve functionally distinct HSC subsets. However, a significant proportion of the cells still lack long-term (LT) reconstituting ability. Therefore, prospective identification of functional HSCs remains crucial to resolve HSC heterogeneity and permit mechanistic analysis of distinct HSC behavior. Here, we used a combination of CD49b and CD229 to subfractionate the phenotypic HSC compartment and identified a CD49b^+^CD229^+^ subset enriched for MPPs with high proliferative activity and short-term reconstituting ability. Our study demonstrates that including additional cell surface markers to the conventional HSC immunophenotype can improve the identification of functional HSCs.

## MATERIALS AND METHODS

### Animal experiments

All animal experiments were approved by the Swedish regional ethical committee, Linköping ethical committee (#882 and #02250-2022). Animals were bred at the Preclinical Laboratory at Karolinska University hospital, Sweden. Young adult (2-5 months) and old (1.5-2 years) females and males on C57BL/6J background were used. In transplantations, Gata-1 eGFP mice (CD45.2),^17^ backcrossed to C57BL/6J (>8 generations), were used as donors. B6.SJL-PtprcaPepcb/BoyCrl and B6.SJL-PtprcaPepcb/BoyJ mice (CD45.1), myeloablated with 10 Gy split over two days, were used as recipients. In primary transplantation, five donor (CD45.2) and 200,000 bone marrow (BM) support (CD45.1) cells were transplanted intravenously. In secondary transplantation, 10 million unfractionated BM cells from primary recipients were each transplanted into 1-2 myeloablated mice. Peripheral blood (PB) was collected periodically from transplanted mice by tail vein bleeding.

The threshold for successful repopulation was based on ≥0.1% CD45.2 and/or eGFP^+^ platelet reconstitution in PB two months post-transplantation. Repopulation of mature blood cells in PB (platelets, erythrocytes, myeloid, B, T, and NK cells) was considered if reconstitution was ≥0.01% and had ≥10 events in the CD45.2 donor gate for any given lineage.^3,15^ These criteria were used to avoid excluding lineage-restricted HSC clones and low-output HSCs, including latent HSCs.^3,6,8,10,15^ Flow cytometric detection limit was determined by assessing false positive events in the CD45.2 donor gates of CD45.1 control mice (n = 12), which was 0.0003%. To determine the number of LT myeloid reconstituted mice, the threshold was set to ≥0.1% myeloid repopulation in PB, 6 months post-transplantation. Using the number of LT myeloid reconstituted mice, HSC frequencies was estimated by limiting dilution analysis through the online tool https://bioinf.wehi.edu.au/software/elda/, calculated with a 95% confidence interval.^18^ These frequencies were used to calculate the HSC frequencies in secondary transplantation by also considering the number of LT myeloid reconstituted mice in secondary transplantation.

### Flow Cytometry experiments

Bone marrow and PB cells were prepared and stained with antibodies (Supplementary Table S1), as previously described.^15^ Cell cycle and cell proliferation analyses were performed according to manufacturer’s protocols (BD Biosciences) and previous studies.^15^ Flow cytometry experiments were performed on FACSAria™ Fusion, LSR Fortessa™ and FACSymphony™ A5 (BD Biosciences). Gates were set based on fluorescent minus one controls, backgating, or negative cell populations. Purity analysis was performed in all cell sorting experiments with a mean purity of 88% (±9%). Post-acquisition analyses were performed on FlowJo software v10 (BD Biosciences).

### In vitro Assays

Combined myeloid and B cell lineage differentiation potential of single sorted cells was assessed *in vitro* using the OP9 co-culture assay.^15^ Cultures were analyzed by flow cytometry three weeks post-sort.

### Statistical Analysis

Statistical analyses were performed using GraphPad v9.4.1 for Mac OS. Non-parametric analysis was performed using Kruskal Wallis with Dunn’s multiple comparison test or Mann-Whitney test. Data are represented as mean ± SD and *p*-values are shown.

## RESULTS AND DISCUSSION

The most primitive HSCs reside within the phenotypic LSKCD48^−^CD34^−^CD150^hi^ (CD150^hi^) compartment in the BM.^2,8,19,20^ CD49b and CD229 have been suggested to subfractionate the functionally heterogeneous HSC population into LT reconstituting HSCs with myeloid-or lymphoid-bias, and MPPs with limited self-renewal ability.^15,16^ Here, we explored whether the combination of CD49b and CD229 can purify functional HSCs, and HSCs with distinct behaviors. CD49b and CD229 subfractionation revealed four immunophenotypic subtypes: CD49b^−^CD229^−^, CD49b^−^CD229^+^, CD49b^+^CD229^−^, and CD49b^+^CD229^+^, which exhibited small but significant differences between females and males (Fig. 1A, 1B; Supplementary Fig. S1A, S1B). We first analyzed whether cycling differences could functionally distinguish the subtypes. All fractions were predominantly found in G0 phase. However, CD49b^−^CD229^−^ and CD49b^−^ CD229^+^ were most quiescent, with the lowest frequency of cells in G1 (Fig. 1C, 1D). Furthermore, BrdU analysis showed that CD49b^−^CD229^−^ and CD49b^−^CD229^+^ were the least proliferative subsets (Fig. 1E, 1F), suggesting that CD49b^−^ subsets represent the most dormant cells regardless of CD229 expression. Females and males exhibited similar cell cycle and proliferation properties across all subtypes (Supplementary Fig. S1C, S1D). Therefore, despite differences in population frequencies, functional characteristics do not differ between sexes.

**Figure 1.**
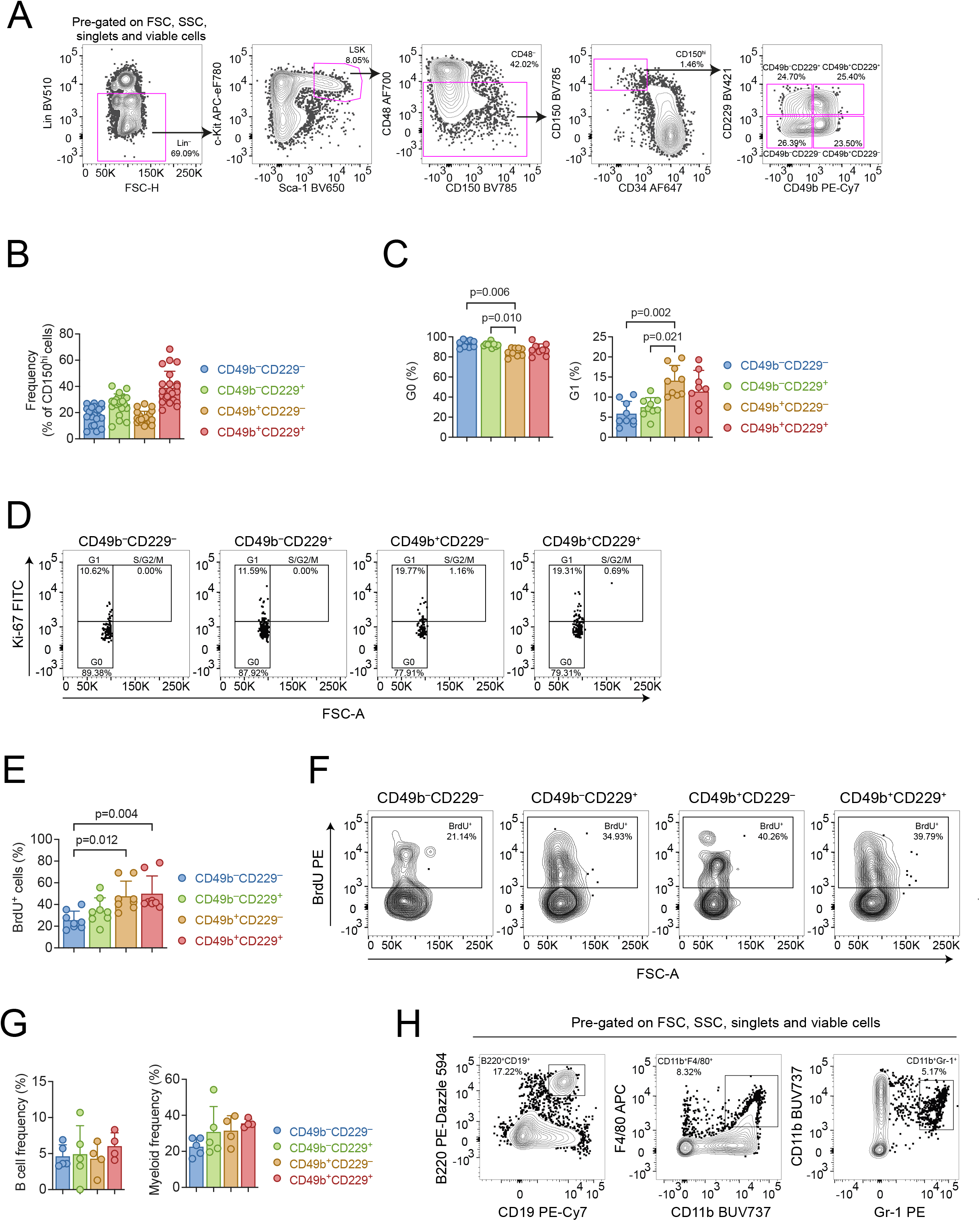
CD49b and CD229 are heterogeneously expressed in the phenotypic HSC compartment. **(A)**: Representative FACS profiles and gating strategy of the phenotypic HSC compartment (Lineage^−^Sca-1^+^c-kit^+^ (LSK) CD48^−^CD34^−^CD150^hi^) and further subfractionation with CD49b and CD229. **(B)**: Frequency of CD49b^−^CD229^−^, CD49b^−^CD229^+^, CD49b^+^CD229^−^, and CD49b^+^CD229^+^ subsets within the phenotypic HSC compartment of young adult mice. n = 22 biological replicates, 9 experiments **(C)**: Cell cycle analysis of CD49b^−^CD229^−^, CD49b^−^ CD229^+^, CD49b^+^CD229^−^, and CD49b^+^CD229^+^ subsets by Ki-67 and DAPI staining in young adult mice. Frequency of cells in G0 (left) and G1 (right) phases are shown. n = 9 biological replicates, 2 experiments. **(D):** Representative FACS profile of cell cycle analysis. **(E):** Cell proliferation analysis by BrdU incorporation in young adult mice. Frequencies of BrdU^+^ CD49b^−^ CD229^−^, CD49b^−^CD229^+^, CD49b^+^CD229^−^, and CD49b^+^CD229^+^ cells are shown. n = 8 biological replicates, 2 experiments. **(F)**: Representative FACS profile of BrdU proliferation analysis. **(G)**: *In vitro* analysis of the B cell and myeloid cell differentiation potential from single cell sorted CD49b^−^CD229^−^, CD49b^−^CD229^+^, CD49b^+^CD229^−^, and CD49b^+^CD229^+^ populations, from young adult mice, using OP9 co-culture assay. Total frequencies of clones with B cells and/or B and myeloid cells (left), and clones containing only myeloid cells (right) are shown. n = 5 biological replicates, 2 experiments. **(H)**: Representative FACS profile of OP9 readout. B cells were defined as CD19^+^B220^+^ and myeloid cells as Gr-1^+^CD11b^+^ and/or F4/80^+^CD11b^+^. Data are represented as mean ± SD. Frequencies of parent gates are shown in FACS plots. Statistical analysis was performed using the Kruskal Wallis test with Dunn’s multiple comparison test in (B-C, F). Abbreviations: FSC, forward scatter; SSC, side scatter; FSC-H, forward scatter height; FSC-A, forward scatter area; BrdU, Bromodeoxyuridine.

Given that both CD49b and CD229 were described to enrich for lymphoid-biased cells,^15,16^ we first investigated the multilineage differentiation ability of CD49b and CD229 fractionated subsets. All four populations could generate B lymphocytes and myeloid cells *in vitro* (Fig. 1G, 1H). We subsequently assessed the multilineage ability *in vivo* by competitive transplantation using limiting numbers from each subtype, with success rates between 73-86% (Supplementary Fig. S1E). The comparable transplantation efficiencies show that the subtypes have similar homing ability. All populations also exhibited similar total blood leukocyte repopulation efficiency, and were capable of multilineage differentiation (Fig. 2A, 2B). Notably, only the CD49b^+^CD229^+^ subtype displayed declining myeloid, platelet, and erythrocyte

**Figure 2.**
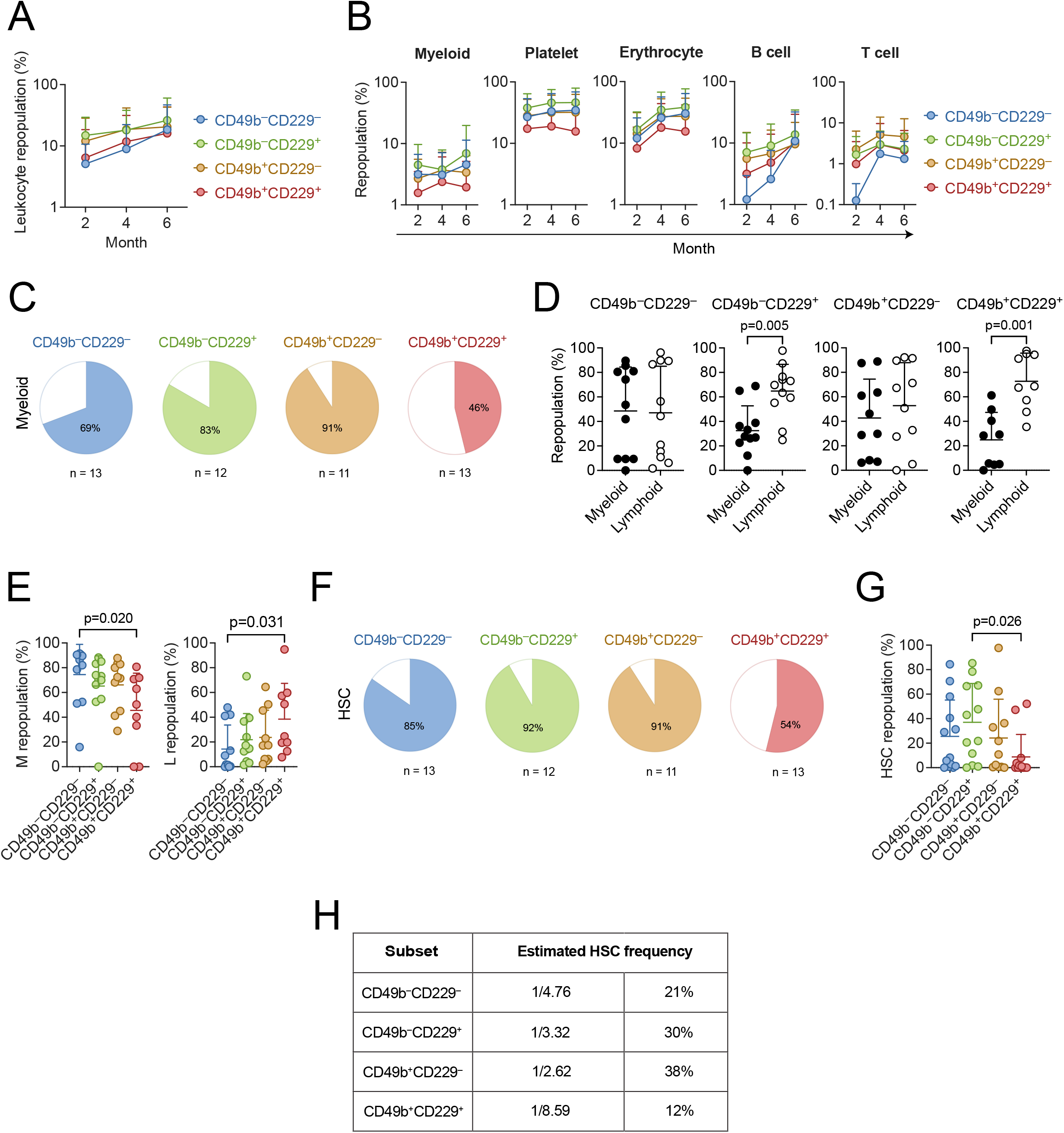
CD49b and CD229 subfractionate the phenotypic HSC compartment into functional HSCs and MPPs with short-term activity. **(A)**: Leukocyte repopulation in the peripheral blood of mice transplanted with 5 cells from CD49b^−^CD229^−^, CD49b^−^CD229^+^, CD49b^+^CD229^−^, and CD49b^+^CD229^+^ populations. **(B)**: Donor-derived myeloid, platelet, erythrocyte, B cell, and T cell frequency in the PB of mice transplanted with 5 cells from CD49b^−^CD229^−^, CD49b^−^CD229^+^, CD49b^+^CD229^−^, and CD49b^+^CD229^+^ populations. **(C)**: Proportion of transplanted mice with ≥0.1% donor-derived myeloid cells in the PB, 6 months post-transplantation. **(D)**: Relative myeloid and lymphoid (B, T, and natural killer cells) repopulation of reconstituted mice in the PB. Mice are selected based on ≥0.1% donor leukocyte (CD45.2^+^) reconstitution in the PB, 6 months post-transplantation. **(E)**: Relative donor-derived myeloid (M, left) and lymphoid (L, right) frequency (L: B, T, and natural killer cells) of reconstituted mice in the BM. Mice are selected based on ≥0.1% donor leukocyte (CD45.2^+^) reconstitution in the BM, 6 months post-transplantation. **(F)**: Proportion of transplanted mice with phenotypic HSC reconstitution. **(G)**: Donor-derived HSC (Lineage^−^Sca-1^+^c-kit^+^Flt-3^−^CD48^−^CD150^+^) frequency in the BM of primary transplanted animals. **(H)**: Estimated HSC frequency based on limiting dilution calculation using the number of mice positively reconstituted in PB myeloid cells, 6 months post-transplantation. n_CD49b_^−^_CD229_– = 13 mice, n_CD49b_^−^_CD229_^+^ = 12 mice, n_CD49b_^+^_CD229_^−^ = 11 mice, and n_CD49b_^+^_CD229_^+^ = 13 = mice, and n_CD49b_ mice, 2 experiments, in (A-C, F-G). n_CD49b_^−^_CD229_– = 11, n_CD49b_^−^_CD229_^+^ = 11, n_CD49b_^+^_CD229_^−^ = 10, and n_CD49b_^+^_CD229_^+^ = 9, 2 experiments, in (D-E) Data are represented as mean ± SD in (A-B, D-E, G). Statistical analysis was performed using the Kruskal Wallis test with Dunn’s multiple comparison test in (A-B and E, G) and the Mann-Whitney test in (D). Abbreviations: M, myeloid; L, lymphoid; HSC, hematopoietic stem cell

reconstitution over time, while other subsets had either increasing, or stable LT repopulation of all blood lineages (Fig. 2B; Supplementary Fig. S2), suggesting reduced reconstituting ability and limited self-renewal potential. Indeed, only 46% of CD49b^+^CD229^+^ transplanted mice could actively produce short-lived myeloid cells six months after transplantation (Fig. 2C). Additionally, CD49b^+^CD229^+^ transplanted mice showed a strong preference for lymphoid cells in the PB, consistent with reduced myeloid output (Fig. 2D). Moreover, the CD49b^+^CD229^+^ subtype generated myeloid cells with the lowest efficiency but potently gave rise to lymphoid cells in the BM (Fig. 2E). These findings imply that CD49b^+^CD229^+^ cells have less HSC activity. In contrast, CD49b^−^CD229^−^, CD49b^−^CD229^+^, and CD49b^+^CD229^−^ subtypes demonstrated robust myeloid reconstitution in the PB and BM, suggesting high self-renewal potential (Fig. 2B-2E). Remarkably, CD49b^−^CD229^+^ cells also showed a strong lymphoid differentiation profile in the PB, indicating that CD49b and CD229 subfractionation can distinguish between HSCs with distinct repopulation characteristics, in addition to differences in self-renewal potential (Fig. 2D; Supplementary Fig. S3).

To further address differences in reconstitution and self-renewal potential, we assessed the ability of CD49b and CD229 fractionated subsets to regenerate stem- and progenitor cells. Only 54% of CD49b^+^CD229^+^ transplanted mice reconstituted phenotypic HSCs, and the repopulation efficiency was significantly lower (Fig. 2F, 2G). These results agreed with a reduced regenerative capacity in CD49b^+^CD229^+^ cells. Furthermore, the number of CD49b^+^CD229^+^ transplanted mice which repopulated lymphoid-primed multipotent progenitors (LMPPs), common lymphoid progenitors (CLPs), granulocyte-monocyte progenitors (GMPs), and megakaryocyte progenitors (MkPs) was markedly reduced compared to other subtypes (Supplementary Fig. S4A). Collectively, these results suggest that the CD49b^+^CD229^+^ subset has reduced HSC activity and enriches for MPPs with short-term repopulating capacity. Therefore, to estimate the frequency of functional HSCs in the CD49b and CD229 fractionated subsets, we used limiting dilution calculation,^18,21^ which predicted the lowest frequency in CD49b^+^CD229^+^ cells, consistent with lower reconstituting ability and reduced HSC activity (Fig. 2H).

To further investigate the self-renewal potential in the CD49b^+^CD229^+^ subset, we selected transplanted mice with phenotypic HSC reconstitution (Fig. 2F, 2G), for secondary transplantation to assess their LT reconstituting ability. All populations could repopulate secondary recipients, however, the CD49b^+^CD229^+^ cells were the least efficient in repopulating both myeloid and lymphoid lineages (Fig. 3A, 3B). Furthermore, the CD49b^+^CD229^+^ group had the fewest animals exhibiting active LT myeloid reconstitution (Fig. 3C). Additionally, the HSC frequency was further reduced in the CD49b^+^CD229^+^ population, in secondary transplantation compared to primary transplantation (Fig. 3D). These findings demonstrate that most CD49b^+^CD229^+^ cells lack extensive self-renewal potential. Conversely, CD49b^−^CD229^−^ and CD49b^−^CD229^+^ subsets maintained a high HSC frequency, consistent with their potent reconstituting ability (Fig. 3B, 3D). The CD49b^+^CD229^−^ subset showed the highest HSC frequency while preferentially repopulating mature lymphoid cells and lymphoid progenitors (LMPP and CLP, Fig. 3B; Supplementary Fig. S4A). We have previously shown that CD49b^+^ HSCs, without CD229 subfractionation, contain LT lymphoid-biased HSCs with an HSC frequency of 11%.^15^ The higher HSC frequency (38%) and preferential lymphoid repopulation in CD49b^+^CD229^−^ cells, suggest that lymphoid-biased HSCs within CD49b^+^ cells can be enriched using CD229 (Fig. 2H, 3B, 3D).

**Figure 3.**
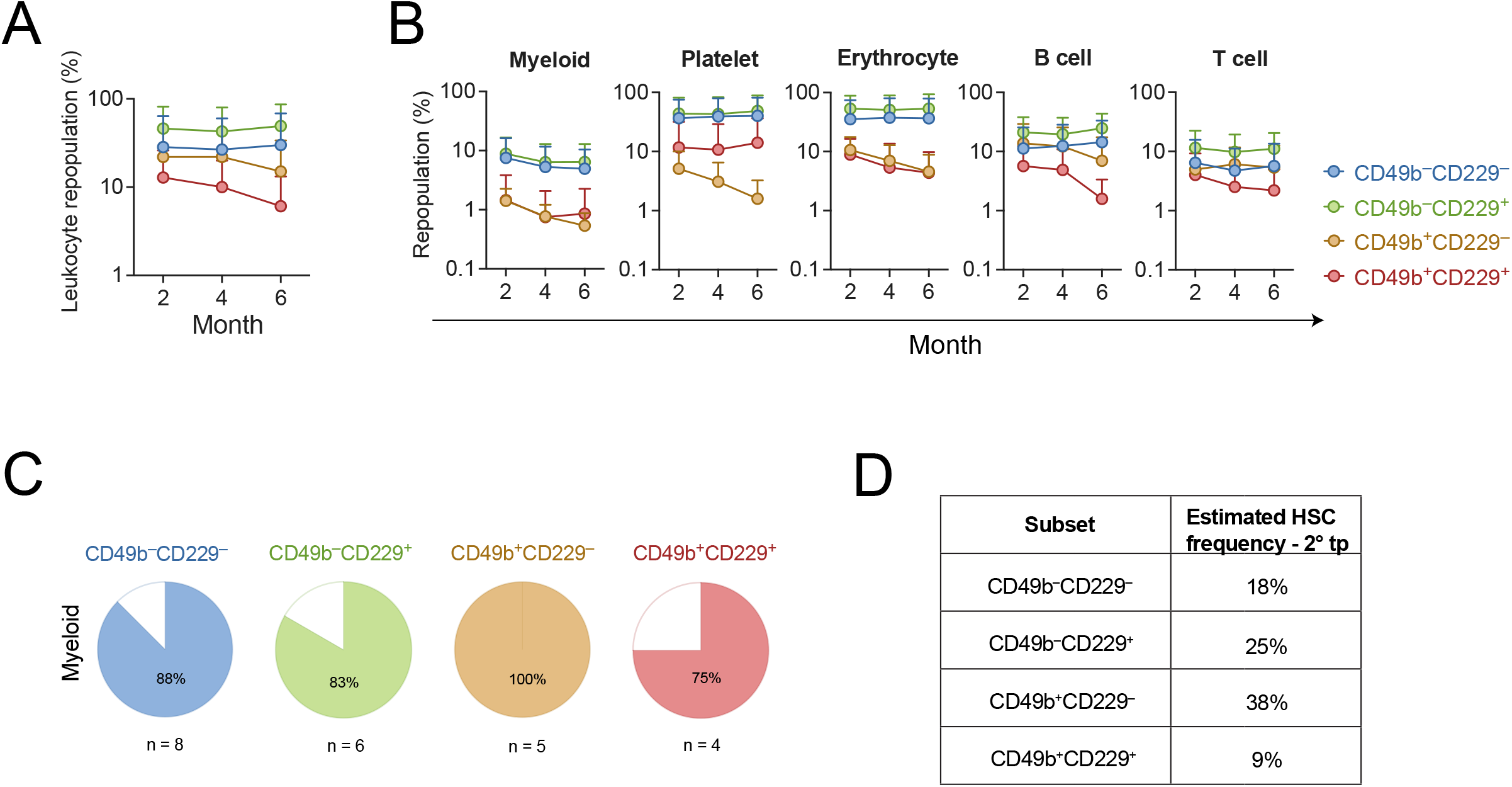
The CD49b^+^CD229^+^ subset lacks extensive long-term reconstituting ability. **(A)**: Leukocyte repopulation in the peripheral blood of secondary transplanted mice from CD49b^−^ CD229^−^, CD49b^−^CD229^+^, CD49b^+^CD229^−^, and CD49b^+^CD229^+^ populations. **(B)**: Donor-derived myeloid, platelet, erythrocyte, B cell, and T cell frequency in the PB of secondary transplanted mice. n_CD49b_^−^_CD229_^−^ = 9 mice, n_CD49b_^−^_CD229_^+^ = 6-7 mice, n_CD49b_^+^_CD229_^−^ = 5 mice, and n_CD49b_^+^_CD229_^+^ = 4-5 mice, 1 experiment, in (A-B). **(C)**: Proportion of secondary transplanted mice with ≥0.1% donor-derived myeloid cells in the PB, 6 months post-transplantation. n_CD49b_^−^_CD229_^−^ = 8 mice, n_CD49b_^−^_CD229_^+^ = 6 mice, n_CD49b_^+^_CD229_^−^ = 5 mice, and n_CD49b_^+^_CD229_^+^ = 4 mice, 1 experiment. **(D)**: Estimated HSC frequency in secondary transplantation, based on limiting dilution calculation from Fig. 2H and the number of mice with active LT myeloid reconstitution from Fig. 3C. Data are represented as mean ± SD in (A-B).

Lymphoid-biased HSCs are suggested to decrease with age.^22^ We therefore assessed the frequency of CD49b and CD229 subfractionated subsets in old mice (Supplementary Fig. S4B). The frequency of CD49b^+^CD229^−^ and CD49b^+^CD229^+^ subsets reduced with age, consistent with an age-related decline, while CD49b^−^CD229^+^ frequency increased and CD49b^−^ CD229^−^ remained unchanged (Supplementary Fig. S4C).

Altogether, our findings demonstrate that LT HSCs are confined to the CD49b^−^CD229^−^, CD49b^−^CD229^+^, and CD49b^+^CD229^−^ subsets. In contrast, the CD49b^+^CD229^+^ subset within the phenotypic HSC compartment predominantly comprise MPPs with high proliferative activity and ST reconstituting ability, and therefore do not represent functional HSCs.

## CONCLUSION

It is recognized that HSCs isolated by immunophenotype contain cells lacking *in vivo* LT reconstituting ability. Therefore, assessment of HSC function is constricted to retrospective analysis through transplantation assays, where HSCs are defined based on functional readouts.^2^ The inability to prospectively identify functional HSCs has limited mechanistic insights into HSC regulation, as molecular analysis of immunophenotypic HSCs reflects a composite of HSCs and MPPs with ST reconstituting potential. Therefore, identifying strategies to isolate functional HSCs remains necessary to enable mechanistic studies, and is of relevance for clinical applications.^23^ In this study, we used the combination of CD49b and CD229 cell surface marker expression within the conventional phenotypic HSC compartment to resolve HSC functional heterogeneity. We demonstrated that the ST reconstituting activity of phenotypic HSCs is predominantly confined to the CD49b^+^CD229^+^ fraction, which contained the lowest HSC frequency and failed to maintain LT hematopoiesis. Furthermore, through segregating CD49b^+^CD229^+^ cells, lymphoid-biased LT HSCs can be enriched in the CD49b^+^CD229^−^ subset. Additionally, our results implicate that the combination of CD49b and CD229 differentiates HSCs with distinct behaviors, which likely can be refined using additional markers. Defining the immunophenotype for diverse HSCs to reveal the full spectrum of HSC behaviors remains important, especially for gene therapies where isolation of distinct HSC subsets may be preferred depending on disease setting.^23^ In conclusion, we demonstrate that functional HSCs with extensive self-renewal ability can be prospectively enriched by discriminating and excluding CD49b^+^CD229^+^ cells containing MPPs with ST reconstituting activity. Our study provides the basis for prospective isolation of HSCs with improved purity and permits more accurate analyses of the molecular mechanisms underlying specific HSC function and behavior.

## Supporting information

Graphical Abstract

## ACKNOWLEDGEMENTS

The authors thank Prof. C. Nerlov (University of Oxford) for providing the Gata-1 eGFP mouse strain and Prof. SE. Jacobsen (Karolinska Institutet) for helpful discussions.

## DATA AVAILABILITY

The data that support the findings of this study are available from the corresponding author, S.L, upon reasonable request.

## REFERENCES

1. Orkin SH, Zon LI. Hematopoiesis: An Evolving Paradigm for Stem Cell Biology. Cell. 2008;132(4):631–644.

2. Wilkinson AC, Igarashi KJ, Nakauchi H. Haematopoietic stem cell self-renewal in vivo and ex vivo. Nat. Rev. Genet. 2020;21(9):541–554.

3. Carrelha J, Meng Y, Kettyle LM, et al. Hierarchically related lineage-restricted fates of multipotent haematopoietic stem cells. Nature. 2018;554(7690):106–111.

4. Sanjuan-Pla A, Macaulay IC, Jensen CT, et al. Platelet-biased stem cells reside at the apex of the haematopoietic stem-cell hierarchy. Nature. 2013;502(7470):232–236.

5. Yamamoto R, Morita Y, Ooehara J, et al. Clonal Analysis Unveils Self-Renewing Lineage-Restricted Progenitors Generated Directly from Hematopoietic Stem Cells. Cell. 2013;154(5):1112–1126.

6. Yamamoto R, Wilkinson AC, Ooehara J, et al. Large-Scale Clonal Analysis Resolves Aging of the Mouse Hematopoietic Stem Cell Compartment. Cell Stem Cell. 2018;22(4):600–607.e4.

7. Müller-Sieburg CE, Cho RH, Thoman M, Adkins B, Sieburg HB. Deterministic regulation of hematopoietic stem cell self-renewal and differentiation. Blood. 2002;100(4):1302–1309.

8. Morita Y, Ema H, Nakauchi H. Heterogeneity and hierarchy within the most primitive hematopoietic stem cell compartment. J. Exp. Med. 2010;207(6):1173–1182.

9. Challen GA, Boles NC, Chambers SM, Goodell MA. Distinct Hematopoietic Stem Cell Subtypes Are Differentially Regulated by TGF-β1. Cell Stem Cell. 2010;6(3):265–278.

10. Dykstra B, Kent D, Bowie M, et al. Long-Term Propagation of Distinct Hematopoietic Differentiation Programs In Vivo. Cell Stem Cell. 2007;1(2):218–229.

11. Haas S, Trumpp A, Milsom MD. Causes and Consequences of Hematopoietic Stem Cell Heterogeneity. Cell Stem Cell. 2018;22(5):627–638.

12. Osawa M, Hanada K, Hamada H, Nakauchi H. Long-term lymphohematopoietic reconstitution by a single CD34-low/negative hematopoietic stem cell. Science. 1996;273(5272):242–245.

13. Adolfsson J, Borge OJ, Bryder D, et al. Upregulation of Flt3 Expression within the Bone Marrow Lin−Sca1+c-kit+ Stem Cell Compartment Is Accompanied by Loss of Self-Renewal Capacity. Immunity. 2001;15(4):659–669.

14. Kiel MJ, Yilmaz ÖH, Iwashita T, et al. SLAM Family Receptors Distinguish Hematopoietic Stem and Progenitor Cells and Reveal Endothelial Niches for Stem Cells. Cell. 2005;121(7):1109–1121.

15. Somuncular E, Hauenstein J, Khalkar P, et al. CD49b identifies functionally and epigenetically distinct subsets of lineage-biased hematopoietic stem cells. Stem Cell Rep. 2022;17(7):1546–1560.

16. Oguro H, Ding L, Morrison SJ. SLAM Family Markers Resolve Functionally Distinct Subpopulations of Hematopoietic Stem Cells and Multipotent Progenitors. Cell Stem Cell. 2013;13(1):102–116.

17. Drissen R, Buza-Vidas N, Woll P, et al. Distinct myeloid progenitor–differentiation pathways identified through single-cell RNA sequencing. Nat. Immunol. 2016;17(6):666–676.

18. Hu Y, Smyth GK. ELDA: Extreme limiting dilution analysis for comparing depleted and enriched populations in stem cell and other assays. J. Immunol. Methods. 2009;347(1–2):70–78.

19. Kent DG, Copley MR, Benz C, et al. Prospective isolation and molecular characterization of hematopoietic stem cells with durable self-renewal potential. Blood. 2009;113(25):6342–6350.

20. Beerman I, Bhattacharya D, Zandi S, et al. Functionally distinct hematopoietic stem cells modulate hematopoietic lineage potential during aging by a mechanism of clonal expansion. Proc. Natl. Acad. Sci. 2010;107(12):5465–5470.

21. Dykstra B, Olthof S, Schreuder J, Ritsema M, Haan G de. Clonal analysis reveals multiple functional defects of aged murine hematopoietic stem cells. J. Exp. Med. 2011;208(13):2691– 2703.

22. Wahlestedt M, Pronk CJ, Bryder D. Concise Review: Hematopoietic Stem Cell Aging and the Prospects for Rejuvenation. STEM CELLS Transl. Med. 2014;4(2):186–194.

23. Epah J, Schäfer R. Implications of hematopoietic stem cells heterogeneity for gene therapies. Gene Ther. 2021;28(9):528–541.

