## Supplementary Figures for "The combination of CD49b and CD229 reveals a subset of multipotent progenitors with short-term activity within the hematopoietic stem cell compartment"

### Figure S1, related to Figure 1-2

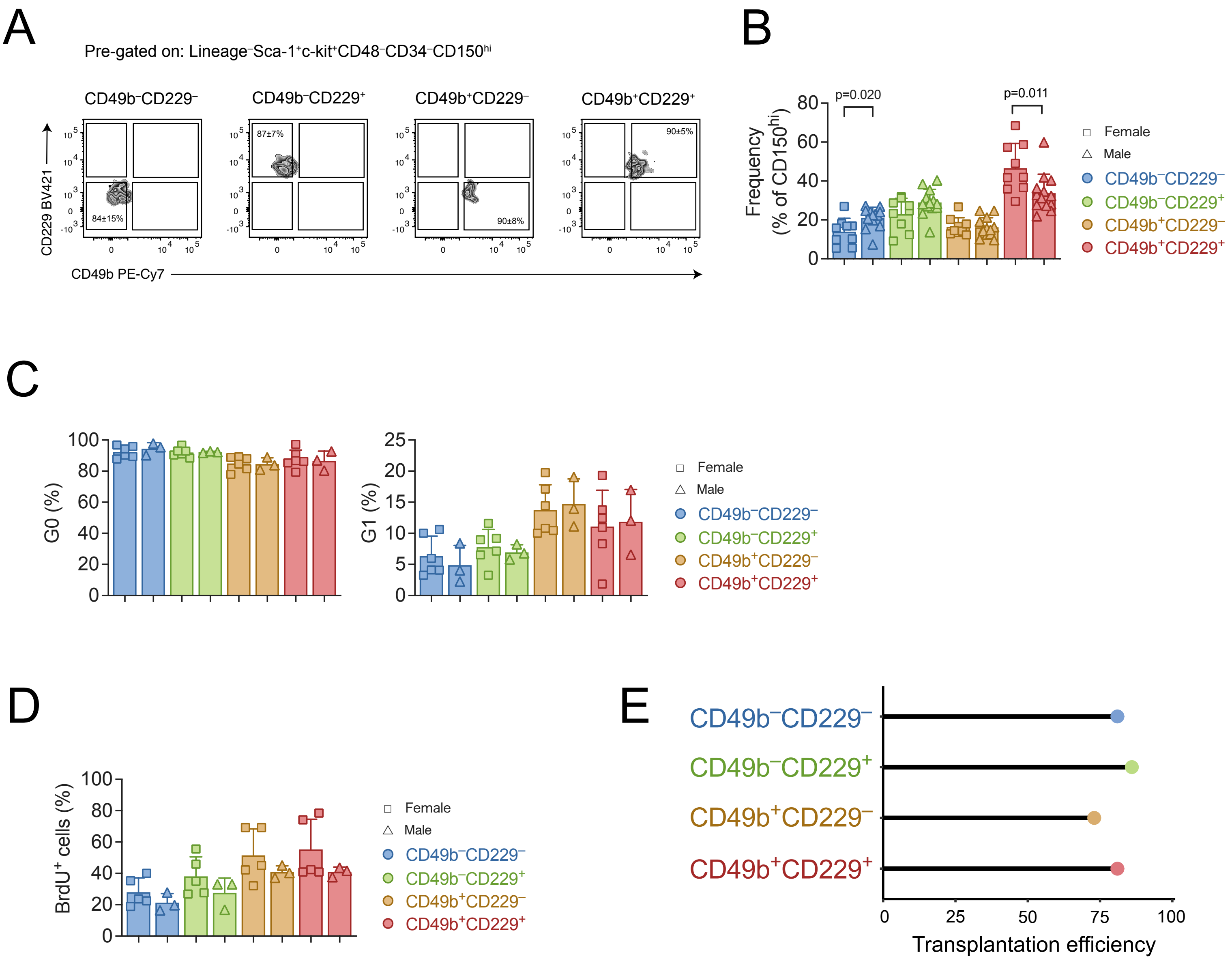

**Supplementary Figure S1, related to Figure 1-2.** CD49b and CD229 subfractionated subsets in female and male mice. **(A):** Sort purity analysis of $CD49b^{-}CD229^{-}$, $CD49b^{-}CD229^{+}$, $CD49b^{+}CD229^{-}$, and $CD49b^{+}CD229^{+}$ populations pre-gated on $Lineage^{-}Sca\text{-}1^{+}c\text{-}kit^{+}CD48^{-}CD34^{-}CD150^{hi}$ ($CD150^{hi}$). Sort purity is represented as mean ± SD for each subset, from 4 experiments **(B)**: Frequency of $CD49b^{-}CD229^{-}$, $CD49b^{-}CD229^{+}$, $CD49b^{+}CD229^{-}$, and $CD49b^{+}CD229^{+}$ subsets within the phenotypic HSC compartment ($CD150^{hi}$) of young adult mice, separated into females and males. n = 9 females, 13 males from 9 experiments **(C)**: Cell cycle analysis of $CD49b^{-}CD229^{-}$, $CD49b^{-}CD229^{+}$, $CD49b^{+}CD229^{-}$, and $CD49b^{+}CD229^{+}$ subsets by Ki-67 and DAPI staining in young adult mice, separated into females and males. Frequency of cells in G0 (left) and G1 (right) phases are shown. n = 6 females, 3 males from 2 experiments. **(D)**: Cell proliferation analysis by BrdU incorporation in young adult mice, separated into females and males. Frequencies of $BrdU^{+}$ $CD49b^{-}CD229^{-}$, $CD49b^{-}CD229^{+}$, $CD49b^{+}CD229^{-}$, and $CD49b^{+}CD229^{+}$ cells are shown. n = 5 females, 3 males from 2 experiments. **(E)**: Transplantation success rate of transplanted $CD49b^{-}CD229^{-}$, $CD49b^{-}CD229^{+}$, $CD49b^{+}CD229^{-}$, and $CD49b^{+}CD229^{+}$ cells.
Statistical analysis was performed with the Mann-Whitney test in (B-D) between female and male groups within the same subset. All data are represented as mean ± SD. Abbreviations: BrdU, Bromodeoxyuridine.

### Figure S2, related to Figure 2

A

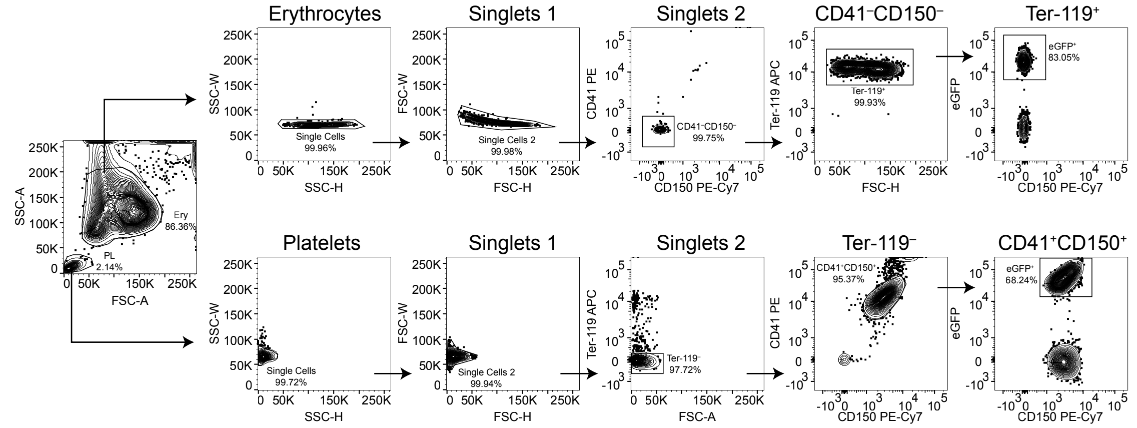


B

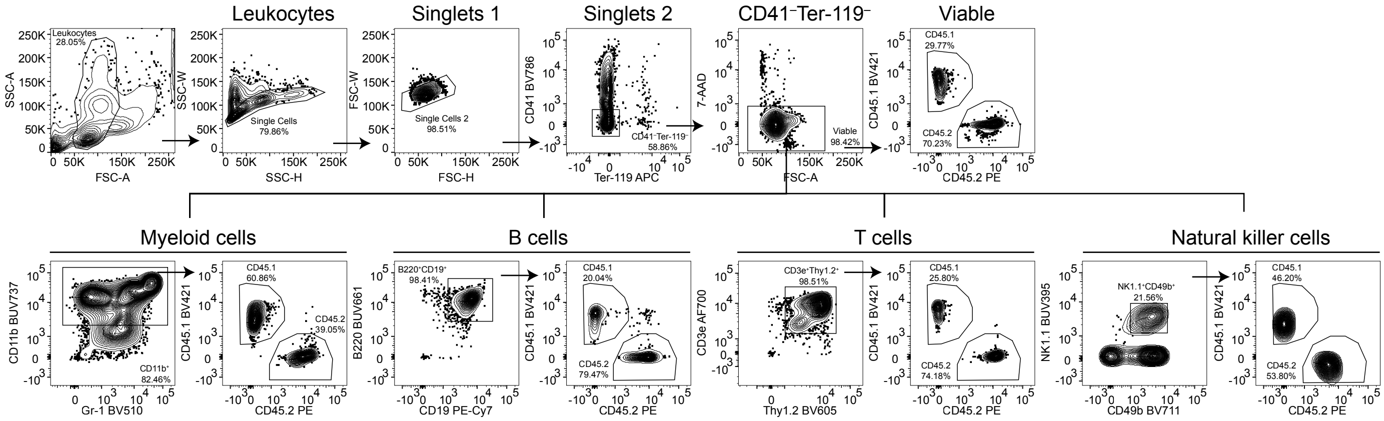

**Supplementary Figure S2, related to Figure 2.** Flow cytometry analysis and gating strategies. **(A)**: Representative FACS profile and gating strategy of platelet and erythrocyte lineages in the peripheral blood. Platelet: *CD41*$^{+}$*CD150*$^{+}$Ter-119$^{-}$; Erythrocyte: *Ter-119*$^{+}$CD41$^{-}$CD150$^{-}$. **(B)**: Representative FACS profile and gating strategy of myeloid, B, T, and natural killer (NK) cell lineages in the peripheral blood. Donor leukocytes: CD45.2$^{+}$; myeloid cell: *CD11b*$^{+}$B220$^{-}$CD19$^{-}$NK1.1$^{-}$CD49b$^{-}$CD3e$^{-}$Thy1.2$^{-}$CD41$^{-}$Ter-119$^{-}$; B cell: *B220*$^{+}$*CD19*$^{+}$ CD11b$^{-}$Gr-1$^{-}$NK1.1$^{-}$CD49b$^{-}$CD3e$^{-}$Thy1.2$^{-}$CD41$^{-}$Ter-119$^{-}$; T cell: *CD3e*$^{+}$*Thy1.2*$^{+}$CD11b$^{-}$Gr-1$^{-}$NK1.1$^{-}$CD49b$^{-}$ B220$^{-}$CD19$^{-}$CD41$^{-}$Ter-119$^{-}$; NK cell: *NK1.1*$^{+}$*CD49b*$^{+}$CD3e$^{-}$Thy1.2$^{-}$CD11b$^{-}$Gr-1$^{-}$B220$^{-}$CD19$^{-}$CD41$^{-}$Ter-119$^{-}$.
Frequency of parent gates are shown in all FACS plots.

### Figure S3, related to Figure 2

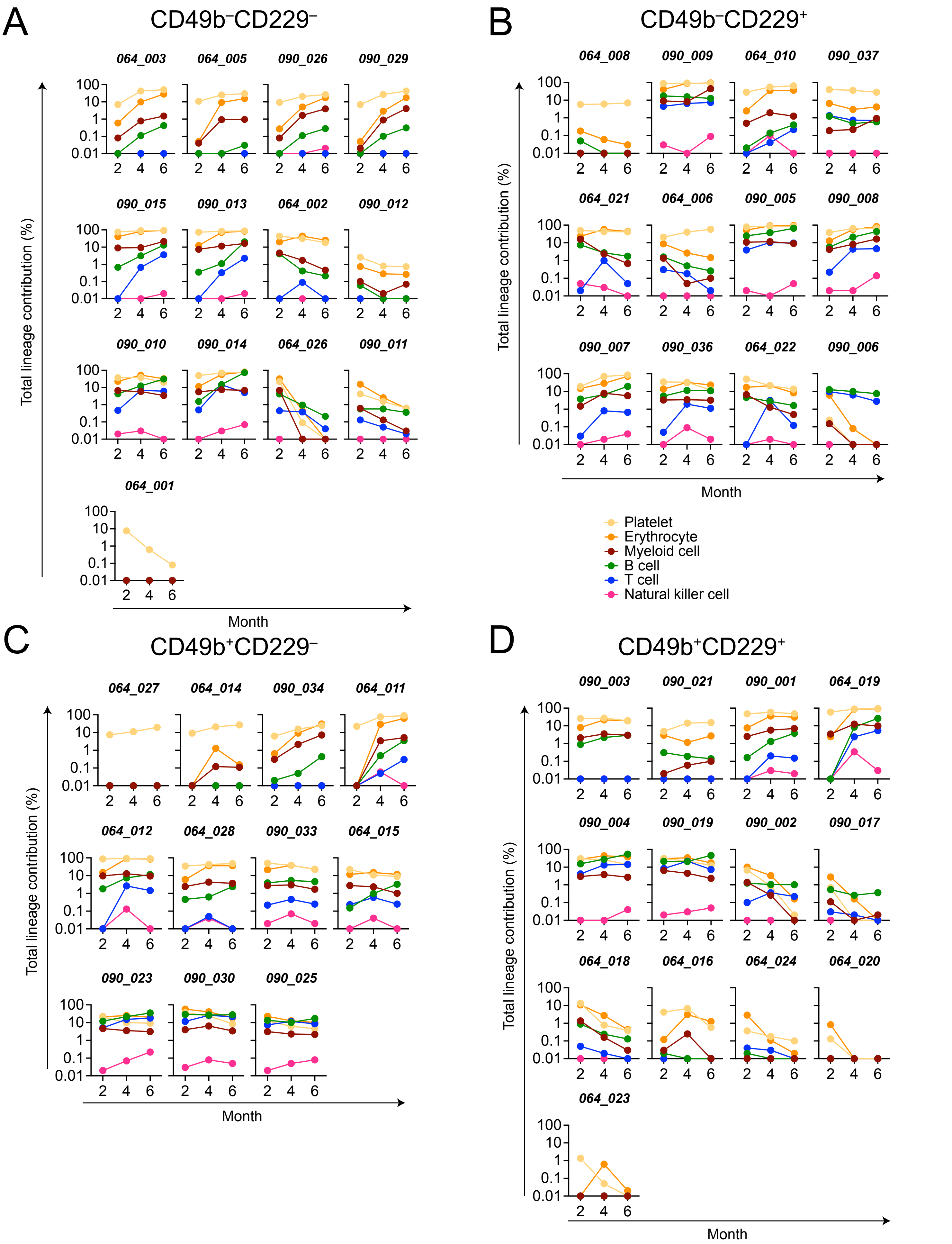

**Supplementary Figure S3, related to Figure 2.** Blood repopulation patterns of primary transplanted mice. **(A-D)**: Frequency of mature lineage repopulation in peripheral blood 2-, 4-, and 6-months post-transplantation. Individual mice, each transplanted with 5 cells from the $CD49b^{-}CD229^{-}$subset in (A), $CD49b^{-}CD229^{+}$ in (B), $CD49b^{+}CD229^{-}$ in (C), and $CD49b^{+}CD229^{+}$ in (D) are shown. Data are represented as mean ± SD.

### Figure S4, related to Figure 2

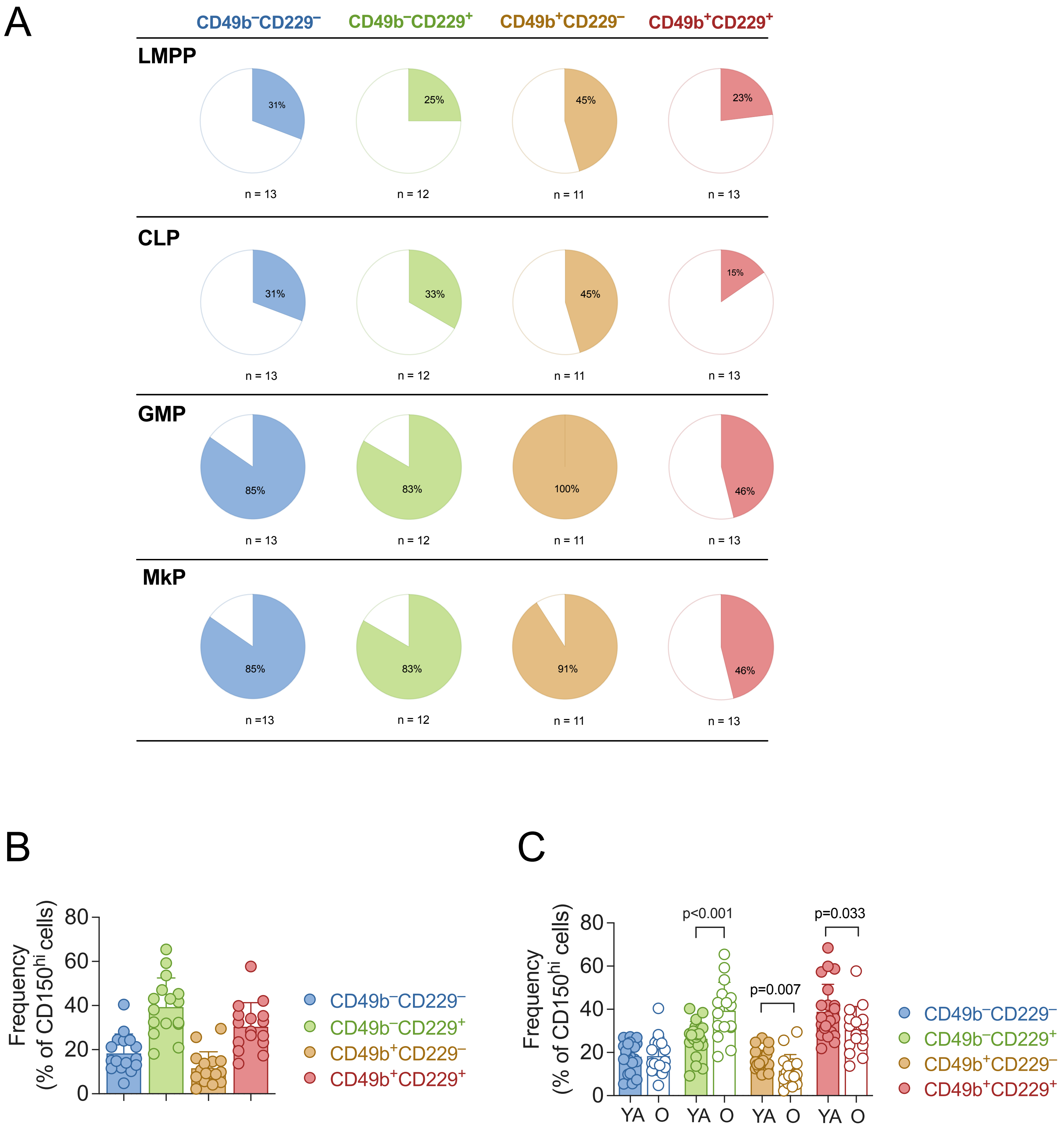

**Supplementary Figure S4, related to Figure 2.** Repopulation of the bone marrow progenitor compartment and CD49b and CD229 subfractionation in aging. **(A)**: Proportion of mice with LMPP, CLP, GMP, and MkP repopulation 6 months post-transplantation. $N_{CD49b^{-}CD229^{-}}$ = 13 mice, $n_{CD49b^{-}CD229^{+}}$ = 12 mice, $n_{CD49b^{+}CD229^{-}}$ = 11 mice, and $n_{CD49b^{+}CD229^{+}}$ = 13 mice, 2 experiments. **(B)**: Frequency of $CD49b^{-}CD229^{-}$, $CD49b^{-}CD229^{+}$, $CD49b^{+}CD229^{-}$, and $CD49b^{+}CD229^{+}$ subsets within the phenotypic HSC compartment ($Lineage^{-}Sca\text{-}1^{+}c\text{-}kit^{+}CD48^{-}CD34^{-}CD150^{hi}$) of old mice. n = 16 biological replicates, 10 experiments. **(C)**: Comparison of the frequency of $CD49b^{-}CD229^{-}$, $CD49b^{-}CD229^{+}$, $CD49b^{+}CD229^{-}$, and $CD49b^{+}CD229^{+}$ subsets in young adult and old mice from Fig. 1B and Fig. S4B.

Statistical analysis was performed with the Mann-Whitney test. Data are represented as mean ± SD in (B-C). Abbreviations: LMPP (Lymphoid-primed multipotent progenitor): $Lineage^{-}Sca\text{-}1^{+}c\text{-}kit^{+}Flt\text{-}3^{hi}$; CLP (Common lymphoid progenitor): $Lineage^{-}B220^{low}Sca\text{-}1^{low}c\text{-}kit^{low}Flt\text{-}3^{hi}IL\text{-}7Ra^{+}$; MkP (Megakaryocyte progenitor): $Lineage^{-}Sca\text{-}1^{-}c\text{-}kit^{+}$ (LK) $CD150^{+}CD41^{+}$; GMP (Granulocyte-monocyte progenitor): LK $CD41^{-}CD150^{-}CD16/32^{+}$; YA, young adult; O, old.

**Supplementary Table S1. Antibody list**

| Antigen | Clone | Supplier | Order number |
|---|---|---|---|
| **Donor and recipient** | | | |
| Anti-mouse CD45.2 (Ly5.2) PE | 104 | BioLegend | 109808 |
| Anti-mouse CD45.1 (Ly5.1) BV421 | A20 | BD Biosciences | 563983 |
| Anti-mouse CD45.1 (Ly5.1) BUV395 | A20 | BD Biosciences | 565212 |
| **Live/Dead, cell proliferation and cell cycle** | | | |
| 7-AAD | | BD Biosciences | 559925 |
| DAPI | | ThermoFisher Scientific | D3571 |
| BrdU PE | | BD Biosciences | 556029 |
| Anti-mouse/Human Ki-67 PE | | BD Biosciences | 567719 |
| Ki-67 FITC | | BD Biosciences | 556026 |
| **Hematopoietic cells** | | | |
| Anti-mouse CD19 PE-Cy7 | 6D5 | BioLegend | 115520 |
| Anti-mouse CD45R/B220 BUV395 | RA3-6B2 | BD Biosciences | 563793 |
| Anti-mouse CD45R/B220 BUV661 | RA3-6B2 | BD Biosciences | 565077 |
| Anti-mouse/Human CD45R/B220 PE-Dazzle 594 | RA3-6B2 | BioLegend | 103258 |
| Anti-mouse/Human CD45R/B220 BV510 | RA3-6B2 | BioLegend | 103248 |
| Anti-mouse CD11b (Mac-1) BUV395 | M1/70 | BD Biosciences | 563553 |
| Anti-mouse CD11b (Mac-1) BUV737 | M1/70 | BD Biosciences | 564443 |
| Anti-mouse/Human CD11b (Mac-1) BV510 | M1/70 | BioLegend | 101263 |
| Anti-mouse F4/80 APC | BM8 | BioLegend | 123116 |
| Anti-mouse F4/80 APC | BM8 | ThermoFisher Scientific | 17-4801-82 |
| Anti-mouse Gr-1 BV510 | RB6-8C5 | BioLegend | 108437 |
| Anti-mouse Gr-1 BUV395 | RB6-8C5 | BD Biosciences | 563849 |
| Anti-mouse Gr-1 (Ly-6G/Ly-6C) PE | RB6-8C5 | BD Biosciences | 553128 |
| Anti-mouse Thy1.2 BV605 | 53-2.1 | BioLegend | 140318 |
| Anti-mouse CD3e AF700 | 500A2 | BD Biosciences | 557984 |

| Antigen | Clone | Supplier | Order number |
|---|---|---|---|
| Anti-mouse CD3e BV510 | 145-2C11 | BioLegend | 100353 |
| Anti-mouse CD3e BUV395 | 1451-2C11 | BD Biosciences | 563565 |
| Anti-mouse CD4 BUV395 | RM4-5 | BD Biosciences | 740208 |
| Anti-mouse CD5 BV510 | 53-7.3 | BioLegend | 100627 |
| Anti-mouse CD5 BUV395 | 53-7.3 | BD Biosciences | 740206 |
| Anti-mouse CD8a BUV395 | 53-6.7 | BD Biosciences | 563786 |
| Anti-mouse NK1.1 BUV395 | PK136 | BD Biosciences | 564144 |
| Anti-mouse CD41 BV786 | MWReg30 | BD Biosciences | 740903 |
| Anti-mouse CD41 PE | eBioMWReg30 | ThermoFisher Scientific | 12-0411-83 |
| Anti-mouse Ter-119 BV510 | TER-119 | BD Biosciences | 563995 |
| Anti-mouse Ter-119 BV650 | TER-119 | BD Biosciences | 747739 |
| Anti-mouse Ter-119 BUV395 | TER-119 | BD Biosciences | 563827 |
| Anti-mouse Ter-119 APC | TER-119 | Proteintech | APC-65149 |
| Anti-mouse CD229 (Ly9) Biotin | Ly9ab3 | BioLegend | 122903 |
| Anti-mouse/Human/Rat Streptavidin BV421 | | BioLegend | 405226 |
| Anti-mouse CD229 (Ly9) BV421 | Ly9.7.144 | BD Biosciences | 748251 |
| Anti-mouse CD229 (Ly9) BUV805 | Ly9.7.144 | BD Biosciences | 749390 |
| Anti-mouse CD49b PE-Cy7 | HMα2 | BioLegend | 103518 |
| Anti-mouse CD49b AF647 | HMα2 | BioLegend | 103511 |
| Anti-mouse CD49b BV711 | HMα2 | BD Biosciences | 740704 |
| Anti-mouse CD150 (SLAM) BV785 | TC15-12F12.2 | BioLegend | 115937 |
| Anti-mouse CD150 (SLAM) PE-Cy7 | TC15-12F12.2 | BioLegend | 115914 |
| Anti-mouse CD34 FITC | RAM34 | ThermoFisher Scientific | 11-0341-85 |
| Anti-mouse CD34 AF647 | RAM34 | BD Biosciences | 560230 |
| Anti-mouse CD48 AF700 | HM48-1 | BioLegend | 103426 |
| Anti-mouse CD48 APC | HM48-1 | BioLegend | 103412 |
| Anti-mouse CD117 (c-kit) APC-eF780 | 2B8 | ThermoFisher Scientific | 47-1171-82 |
| Anti-mouse Sca-1 (Ly-6A/E) BV605 | D7 | BioLegend | 108134 |
| Anti-mouse Sca-1 (Ly-6A/E) BV650 | D7 | BD Biosciences | 740450 |

| Antigen | Clone | Supplier | Order number |
|---|---|---|---|
| Anti-mouse CD105 (Endoglin) BV650 | MJ7/18 | BD Biosciences | 740609 |
| Anti-mouse CD135 (Flt-3) PE | A2F10 | BioLegend | 135306 |
| Anti-mouse CD135 (Flt-3) BV421 | A2F10.1 | BD Biosciences | 562898 |
| Anti-mouse CD127 (IL7-Ra) BV711 | A7R34 | BioLegend | 135035 |
| Anti-mouse CD16/32 AF700 | 93 | ThermoFisher Scientific | 56-0161-82 |
| Anti-mouse CD16/32 BUV737 | AB93 | BD Biosciences | 751697 |
| Anti-mouse CD16/32 (Fc-block) Purified | 2.4G2 | BD Biosciences | 553142 |
