## Supplementary figures and images for "The combination of CD49b and CD229 reveals a subset of multipotent progenitors with short-term activity within the hematopoietic stem cell compartment"

### Graphical Abstract

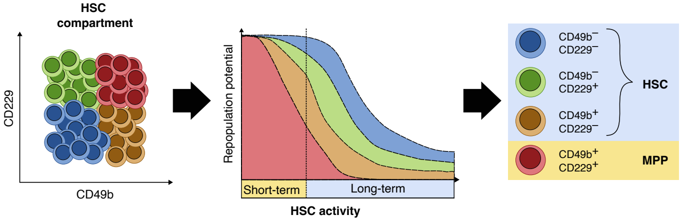
